# SHC-3: a previously unidentified *C. elegans* Shc family member functions in the insulin-like signaling pathway to enhance survival during L1 arrest

**DOI:** 10.1101/2023.11.22.568309

**Authors:** Victoria L. León Guerrero, Mercedes Di Bernardo, W. Rod Hardy, Jacob C. Sutoski, Lesley T. MacNeil

## Abstract

Shc proteins function in many different signaling pathways where they mediate phosphorylation-dependent protein-protein interactions. These proteins are characterized by the presence of two phosphotyrosine-binding domains, an N-terminal PTB and a C-terminal SH2. We describe a previously unrecognized *C. elegans* Shc gene, *shc-3* and characterize its role of in stress response. Both *shc-3* and *shc-1* are required for long-term survival in L1 arrest, however, they do not act redundantly but rather play distinct roles in this process. SHC-3 function in survival during L1 arrest is DAF-16-dependent, demonstrating that like SHC-1, SHC-3 functions in the Insulin-like signaling pathway. In the absence of SHC-3, nuclear entry and exit are slowed suggesting that SHC-3 is required for rapid changes in DAF-16 signaling.

## Introduction

The regulation of protein-protein interactions by phosphorylation allows rapid transmission of signals with tight control over the timing and length of activity. Proteins that bind specific phosphorylated motifs therefore play critical roles in many signaling pathways. Among these proteins are the Shc (Src homologous and collagen) family proteins that are integral to many cellular pathways where they link activated receptors to downstream effectors.

Shc proteins are found in evolutionarily diverse organisms where they regulate a number of different biological processes, including stress response. They have no enzymatic activity but instead function as intracellular signaling scaffolds, regulating signal-dependent complex formation in part through PTB and SH2 phosphotyrosine-binding domains. The N-terminal PTB domain binds motifs matching the θ-X-N-X-X-pY consensus (θ = large hydrophobic amino acid), and a C-terminal SH2 domain that binds motifs matching the consensus, pY-θ-X-θ. This domain organization is unique to members of the Shc family. The central region, CH1 (collagen homologous), exhibits limited sequence similarity between Shc proteins of different species. In mammalian ShcA, this region contains two conserved YXN motifs that, when phosphorylated, recruit Grb2 (Rozakis-Adcock *et al*. 1992). The first of these motifs, YYN (Y239/240 in hShcA) is conserved in *Drosophila* Shc proteins but absent in *C. elegans* Shc proteins while the second motif (Y317 in hShcA) is absent from both *Drosophila* and *C. elegans* Shc proteins, suggesting that the role of Grb2 in Shc signaling may not be conserved in these species.

Two *shc* genes were previously reported in *C. elegans*, *shc-1* and *shc-2 (Luzi et al. 2000)*. Little is known about the function of *shc-2*, but *shc-1* functions in stress response pathways, including the Insulin-like signaling (IIS) and MAPK pathways, to regulate lifespan, pathogen resistance, dauer formation and oxidative stress resistance (Neumann-Haefelin *et al*. 2008). Via its PTB domain, mammalian ShcA binds activated Insulin receptors (IR or IGF-1). This binding leads to phosphorylation of ShcA, preferentially on Y317, and subsequent recruitment of Grb2; Grb2 in turn recruits SOS and activates the ras/MAPK pathway. While *C. elegans* SHC-1 both promotes MAPK signaling and binds the DAF-2 receptor, this is unlikely to occur via the same mechanism because SHC-1 lacks the conserved Grb2 binding sites. Further, SHC-1 is proposed to act as a negative regulator of DAF-2 but as a positive regulator of JNK-MAPK signaling, suggesting the two roles are independent (Neumann-Haefelin *et al*. 2008; Mizuno *et al*. 2008). SHC-1 acts as a scaffold bridging MEK-1 and MLK-1, promoting the phosphorylation of MEK-1 and subsequent activation of the JNK-like MAPK KGB-1. SHC-1 functions with MEK-1 and MLK-1 to regulate oxidative stress response, heavy metal resistance, dauer entry and axon regeneration (Mizuno *et al*. 2008; Hisamoto *et al*. 2016; Dogra *et al*. 2022).

While vertebrates have four Shc genes (Shc1/ShcA, Shc2/ShcB, Shc3/ShcC and Shc4/ShcD), most invertebrates have a single Shc gene that is most similar to ShcA, suggesting that ShcA is the ancestral Shc gene and that this gene family was expanded in the vertebrate lineage. The *C. elegans* genome contains three genes that encode proteins with Shc-like domain structure, SHC-1, SHC-2 (Luzi *et al*. 2000), and a previously uncharacterized protein, K11E4.2, that we have named SHC-3. Here we report the identification and characterization of the *C. elegans* Shc family gene, K11E4.2/*shc-3*. Animals lacking *shc-3* show reduced survival in L1 arrest and are more sensitive to pathogen exposure. We find that, like SHC-1, SHC-3 functions in the IIS pathway but that the two proteins do not function redundantly but instead play distinct roles in regulating IlS.

## Results

### K11E4.2 encodes a SHC-family protein

By examining SH2 domain-containing proteins in *C. elegans*, we identified a previously uncharacterized Shc family protein, K11E4.2, that we have named SHC-3. This protein contains the characteristic Shc-family domain structure, an N-terminal PTB domain and a C-terminal SH2 domain (**Figure 1A**). SHC-3 was not previously identified as a SHC protein because the N-terminal domain was annotated as a PH-superfamily domain rather than a PTB domain. PH domains share the same basic structural fold as PTB domains but differ in function in that PH domains interact with phospholipids, while PTB domains bind phosphotyrosine-containing motifs. Intriguingly, the ShcA PTB domain can interact with both phosphotyrosine and phospholipid (Ravichandran *et al*. 1997), a feature that may explain sequence similarity of the Shc PTB domain and PH domains.

**Figure 1.**
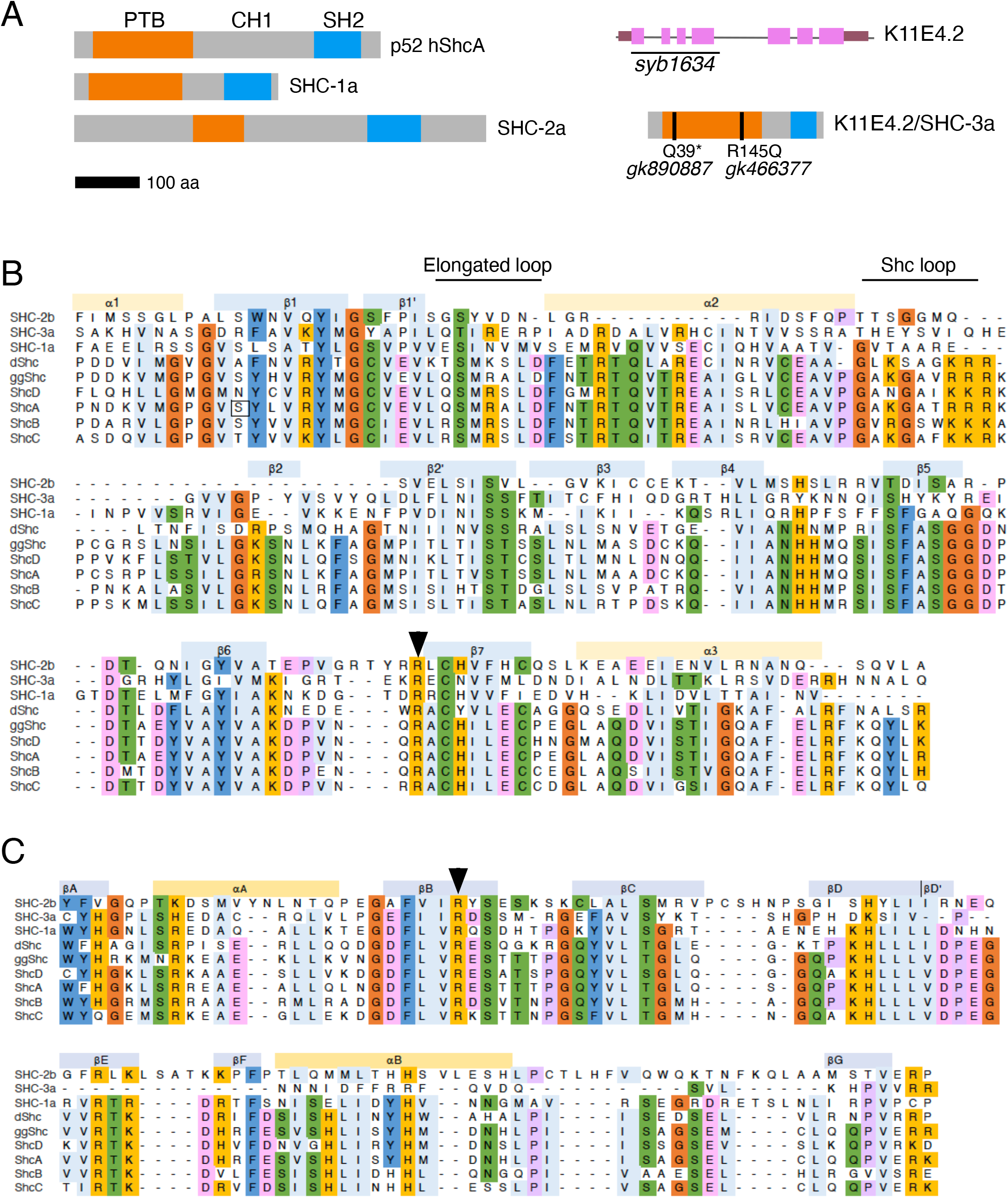
K11E4.2 encodes a Shc-like protein. (A) Domain organization of *C. elegans* SHC proteins. K11E4.2 alleles are shown. (B) Alignment of *C. elegans*, *Drosophila melanogaster* (d), *Galus galus* (gg) and Human (h) Shc PTB domains. Structural features of human ShcA are shown above the alignment. The elongated loop and the Shc loop, characteristic of Shc proteins are indicated. Arrowhead highlights the conserved arginine required for phosphotyrosine binding. (C) Alignment of Shc SH2 domains. Arrowhead indicates conserved arginine required for phosphotyrosine binding.

PTB domains are variable at the amino acid level. Based on specific structural features, they can be categorized as Shc-like, IRS-like, or Dab-like (Zhou *et al*. 1995; Eck *et al*. 1996; Farooq *et al*. 1999; Yun *et al*. 2003; Uhlik *et al*. 2005). K11E4.2 has features of the Shc PTB domain, including the conserved Arginine (R175 of hShcA, R145 in SHC-3) required for phosphotyrosine binding, an elongated loop between the PTB β1 and α2 that is distinct from Dab-like PTB domains, and the ‘Shc loop’, an approximately 20 amino acid loop between α2 and β2 that is absent in IRS1 and DAB PTB domains (Zhou *et al*. 1995; Eck *et al*. 1996)(**Figure 1B Supplemental Figure 1**). These structural features are conserved across *Caenorhabditis* K11E4.2 orthologs (**Supplemental Figure 2**), supporting the identification of the N-terminal domain of K11E4.2 as a Shc-like PTB domain.

A second phospho-tyrosine binding domain, the SH2 domain, lies at the C-terminus of Shc proteins. The N-terminus of the SH2 domain mediates its binding to phosphotyrosine motifs, this region in K11E4.2 is similar to human Shc proteins and the conserved arginine residue required for SH2 domains to bind phosphotyrosine is present in all *Caenorhabditis* SHC-3 sequences examined (**Figure 1C and Supplemental Figure 2**). By contrast, the C-terminus of the SHC-3 SH2 domain is more divergent. Based on sequence alignment and the alpha fold prediction, the βE-βF region is likely absent or shortened relative to the human Shc proteins (**Supplemental Figure 1**). Variability in this region is common in the SH2 domain family, more variability occurs at the C-terminus with deletions and insertions primarily found in the βE-βF and βG loop regions (Kaneko *et al*. 2012). The C-terminus of the *C. elegans* SHC-2 SH2 domain is also divergent, with insertions observed in these regions. Together, this may suggest that the *C. elegans* SHC SH2 domains have different binding specificities or may be differentially regulated through protein-protein interaction or post-translational modification.

One of the most well-characterized functions of mammalian ShcA is the recruitment of Grb2 and associated proteins to tyrosine-phosphorylated transmembrane receptors resulting in the activation of the MAPK pathway. Grb2 binds ShcA through phosphorylated YXN motifs in the CH1 domain of Shc (Rozakis-Adcock *et al*. 1992). Similar to *C. elegans* SHC-1 and SHC-2, SHC-3 has little sequence similarity in the CH1 domain and lacks the YXN motifs that serve as docking sites for GRB2 in mammalian ShcA, suggesting that, like *C. elegans* SHC-1, SHC-3 does not physically associate with SEM-5/Grb2.

### SHC-3 is expressed in the intestine

To examine SHC-3 expression and localization, we generated a SHC-3-GFP translational reporter (*shc-3p::shc-3::GFP*). In contrast to *shc-1,* which is widely expressed (Neumann-Haefelin *et al*. 2008), *shc-3p::shc-3::GFP* expression was restricted to the intestine (**Figure 2A**). Expression was observed from embryogenesis through larval development and into the adult stage. Although both Shc proteins are expressed in the intestine, their subcellular localization is distinct. At 20C, the SHC-3-GFP fusion protein was concentrated at the apical surface but was also observed in the cytoplasm (**Figure 2A**). By contrast, SHC-1 is strongly localized to the nucleus (Neumann-Haefelin *et al*. 2008), suggesting that the two proteins play distinct roles. The concentration of SHC-3, but not SHC-1 at the cell membrane may also suggest that SHC-3 performs the more well-recognized role of Shc proteins, to functions as an adapter for receptor tyrosine kinases.

**Figure 2.**
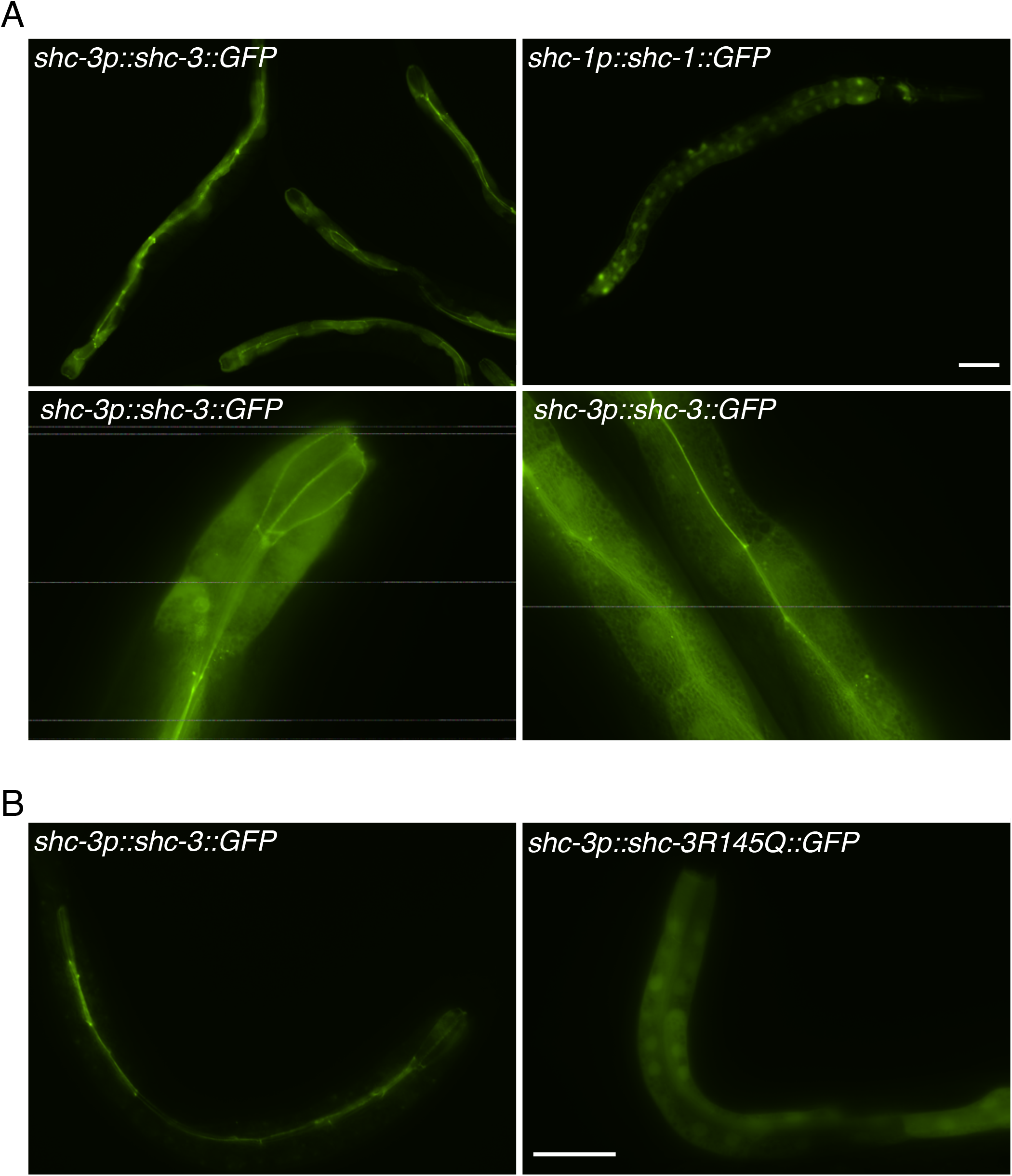
*shc-3* is expressed in the intestine. (A) Intestinal expression with apical localization of *shc-3p::shc-3::GFP* (left), compared to *shc-1p::shc-1::GFP* expression (right). (B) SHC-3::GFP is apically localized in the intestine. (C) Substitution of the conserved R in the SHC-3 PTB domain reduces localization of SHC-3 to the plasma membrane. Scale bar = 100 mm.

Mammalian Shc proteins can be recruited to the plasma membrane through the interaction of the PTB domain with tyrosine-phosphorylated transmembrane proteins or membrane-associated proteins (Ravichandran *et al*. 1997). We reasoned that the inability to bind phosphotyrosine motifs might change the subcellular localization of SHC-3. To test this, we generated a *shc-3* translational fusion carrying a substitution of the conserved arginine (R145Q); the analogous substitution in human ShcA prevents phosphotyrosine binding and reduces membrane localization (Ravichandran *et al*. 1997). SHC-3R154Q::GFP displayed decreased apical localization compared to the wild-type protein, suggesting that, as with ShcA, the PTB domain, likely through phosphotyrosine binding, is required to recruit SHC-3 to the membrane (**Figure 2B**).

### *C. elegans* Shc proteins function in stress response

To examine the function of *shc-3*, we generated a deletion allele that removes most of exons 1-4 and obtained two additional alleles, gk890887 and gk466377, from the million mutation project (Thompson *et al*. 2013). The gk890887 allele produces an early stop (Q39*), while the gk466377 allele produces the R145Q substitution that reduced membrane localization in our transgenic animals (**Figure 1A**). All three mutants appeared superficially wild type in standard growth conditions. We measured growth, reproduction and lifespan in *shc-3* null mutant animals. On an *E. coli* OP50 diet, *shc-3* mutant animals developed normally and produced wild-type brood sizes (**Figure 3A**). Unlike *shc-1* mutants, *shc-3* mutants did not have reduced lifespans, however, in double mutants, loss of *shc-3* suppressed the shortened lifespan of *shc-1* mutants (**Figure 3B**), suggesting that the two proteins may have opposing functions.

**Figure 3.**
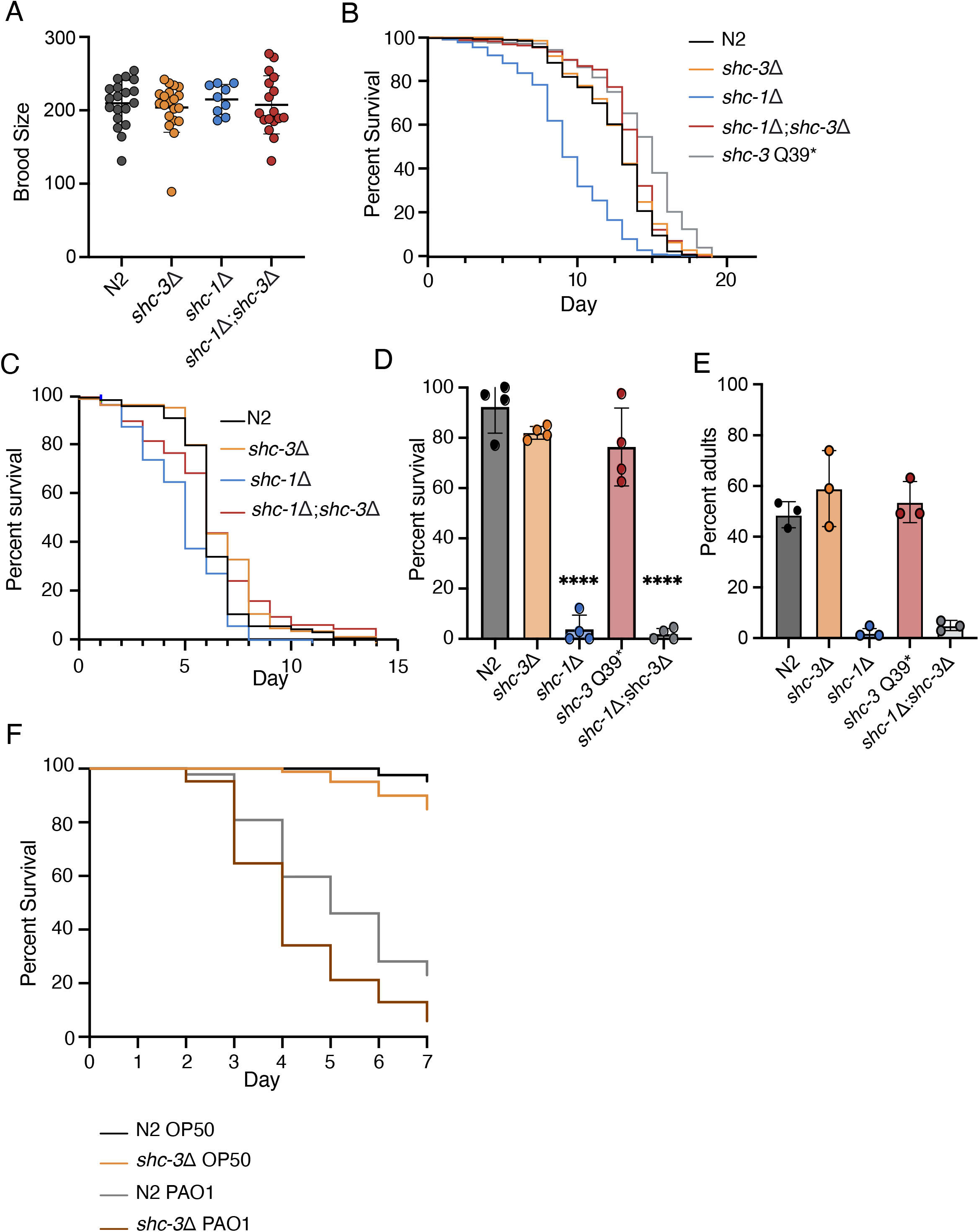
Characterization of *shc-3* mutants. (A) Total brood size of indicated mutants, N2 is used as wild-type (B) Lifespan of Shc mutants. As previously reported, *shc-1* mutants had shortened lifespans (p<0.005). *shc-3* mutation suppressed the shortened lifespan of *shc-1* mutants (p<0.005). Days indicates days of adulthood. Significance calculed by Mantel-Cox test with Bonferroni correction for multiple hypothesis testing. (C) Survival following exposure of adult animals to 4*ℳ*m paraquat. Increased sensitivity of *shc-1* mutants is suppressed by *shc-3* mutation (p<0.005 Wilcoxon test with bonferroni correction) (D) *shc-3* mutants did not have increased sensitivity to CuSO4 (E) or tunicamycin (F) *shc-3* mutants are more sensitive to *P. aeruginosa* PA01 than wild-type animals.

The importance of Shc proteins in stress response is conserved in *C. elegans*. Loss of *shc-1* increases sensitivity to oxidative stress, copper sulfate, tunicamycin, and pathogen exposure (Neumann-Haefelin *et al*. 2008; Mizuno *et al*. 2008). We asked whether *shc-3* was also required for response to these stresses. *shc-3Δ* animals were not more sensitive to oxidative stress induced by paraquat, and loss of *shc-3* suppressed the sensitivity of *shc-1* mutants. *Shc-3Δ* mutants had increased sensitivity to infection by *Pseudomonas aeruginosa* PA01 (**Figure 3D**) but not to CuSO_4_, or tunicamycin (**Figure 3E, F**). Loss of *shc-3* did not enhance or suppress sensitivity to CuSO_4_, or tunicamycin in *shc-1(ok198)* mutants, however, these effects are likely mediated through the hypodermis where only *shc-1* is expressed. *shc-1* expression in the hypodermis is sufficient to rescue the CuSO_4_ sensitivity of *shc-1* mutants (Mizuno *et al*. 2008). Similarly, the tunicamycin sensitivity of *kgb-1* mutants is partially rescued by expression in the hypodermis or muscle but not by intestinal expression of *kgb-1.* As SHC-1 functions upstream of KGB-1, it is likely that loss of SHC-1 in the hypodermis or muscle is responsible for the sensitivity to tunicamycin (Liu *et al*. 2018).

### *Shc-3* is required for long-term survival in L1 arrest

*Shc-1* mutants develop a gonad dysmorphology phenotype following prolonged starvation, suggesting that SHC-1 is required for recovery from starvation (Wolf *et al*. 2014). It may also play a role in recovery from long-term arrest through KGB-1, given *kgb-1* mutants arrested for long periods in L1 are slow to recover from arrest and have decreased lifespans (Roux *et al*. 2016)(Roux *et al*. 2022). We examined the survival of Shc mutants after prolonged L1 arrest. Survival of *shc-1(ok198)* and all three *shc-3* mutants was decreased relative to wild-type animals (**Figure 4A**). The decreased survival of *shc-3* R145Q suggests that the PTB domain is important for this function. While mutants in both Shc genes displayed the same phenotype, there was no enhanced effect when both genes were lost (**Figure 4B**), suggesting that SHC-1 and SHC-3 do not act redundantly but that they do act in the same pathway to regulate survival in L1 arrest.

**Figure 4.**
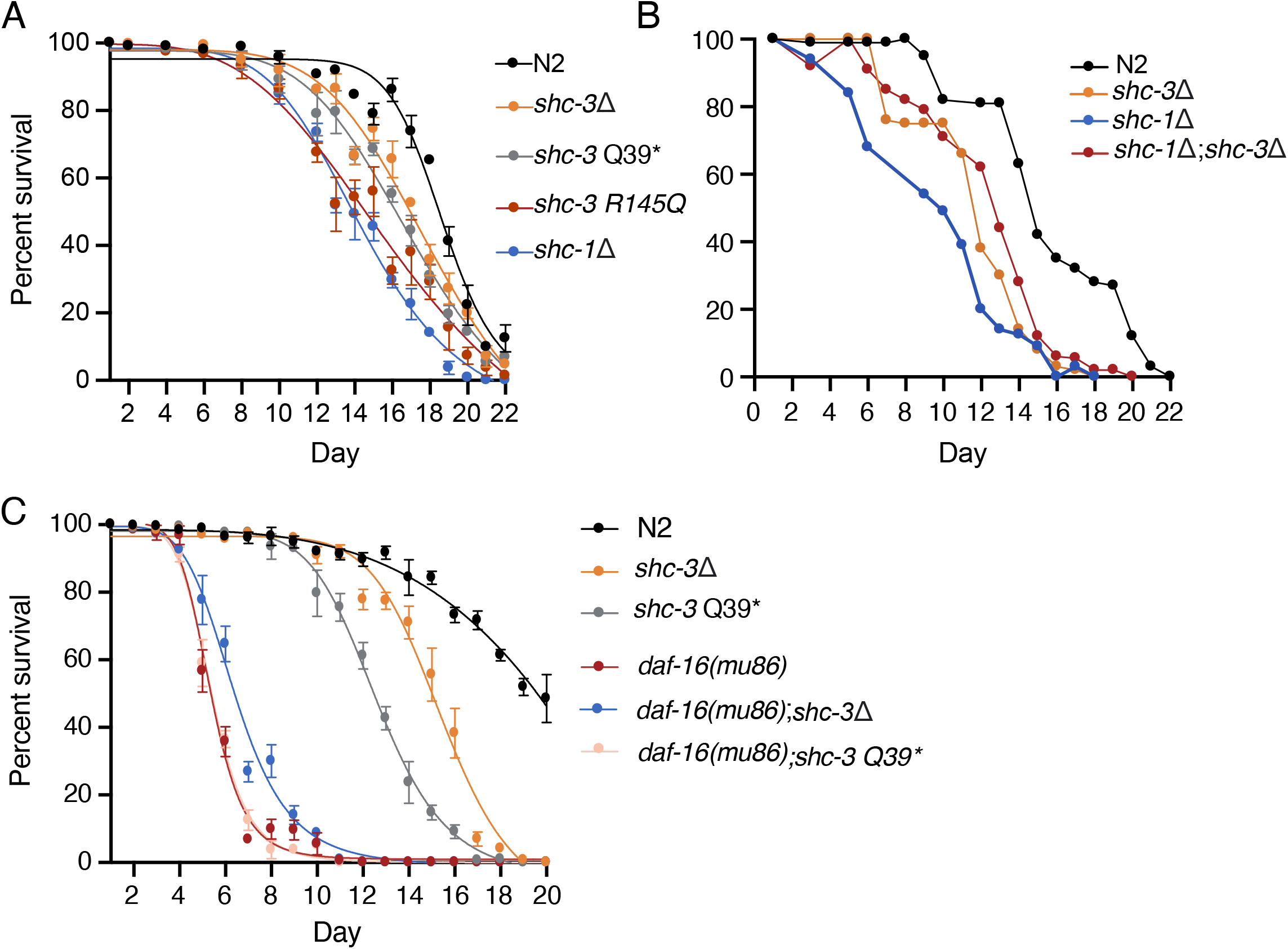
*shc-3* is required for long-term survival in L1 arrest. Animals were hatched in M9 buffer and survival was scored over time (A) *shc-1* and *shc-3* mutants have reduced survival in L1 arrest relative to N2 animals. (B) Reduced survival of *shc-1* and *shc-3* mutants was not further enhanced in double mutants. (C) *shc-3* mutation does not enhance the reduced survival of *daf-16 (mu86)* mutants. Animals were scored in triplicate at each time point.

Insulin-like signaling is a key regulator of survival during L1 arrest in *C. elegans* (Baugh and Sternberg 2006). Loss of the insulin-like receptor *daf-2* or of the PI3K subunit *age-1* increases survival in L1 arrest while loss of their downstream target FOXO/*daf-16* decreases survival (Munoz and Riddle, 2003 (Baugh and Sternberg 2006; Zhang *et al*. 2011)). Two functions have been proposed for SHC-1, as a physical interactor and negative regulator of the Insulin-like receptor DAF-2, and as an activator of JNK signaling; both functions converge on DAF-16 (Neumann-Haefelin *et al*. 2008). To determine if SHC-3 influences L1 arrest via the IIS pathway, we measured survival in L1 arrest of *daf-16* mutants carrying *shc-3Δ* or *shc-3 Q39\** mutations and found that neither allele decreased survival of *daf-16* mutants, suggesting that the function of SHC-3 in L1 arrest is DAF-16-dependent (**Figure 4C**). The similarity of phenotypes between *daf-16* and *shc-3* mutants would suggest that in L1 arrest SHC-3 and SHC-1 function as positive regulators of DAF-16, and therefore negative regulators of IlS.

### SHC-3 regulates DAF-16 nuclear entry and exit

To further understand the relationship between SHC-3 and insulin signaling, we examined the influence of *shc-3* mutation on DAF-16 nuclear localization during stress. Activation of IIS prevents DAF-16 from entering the nucleus and activating its transcriptional targets. Localization of DAF-16::GFP can therefore serve as a proxy for insulin signaling (Henderson and Johnson 2001). In response to a 20 minute heat shock, DAF-16::GFP nuclear localization was decreased in *shc-3* mutant animals relative to wild-type animals (**Figure 5A**). We also examined recovery from heat shock by inducing complete nuclear localization with a long heat shock followed by a 1.5 hour recovery period at 20°C. Under these conditions, more DAF-16::GFP was observed in the nucleus of *shc-3* mutants than controls, suggesting that recovery from heat shock is slowed in these animals (**Figure 5B&C**). The delayed nuclear entry and exit of DAF-16 in *shc-3* mutants suggests that SHC-3 is required to mediate rapid changes in insulin signaling.

**Figure 5.**
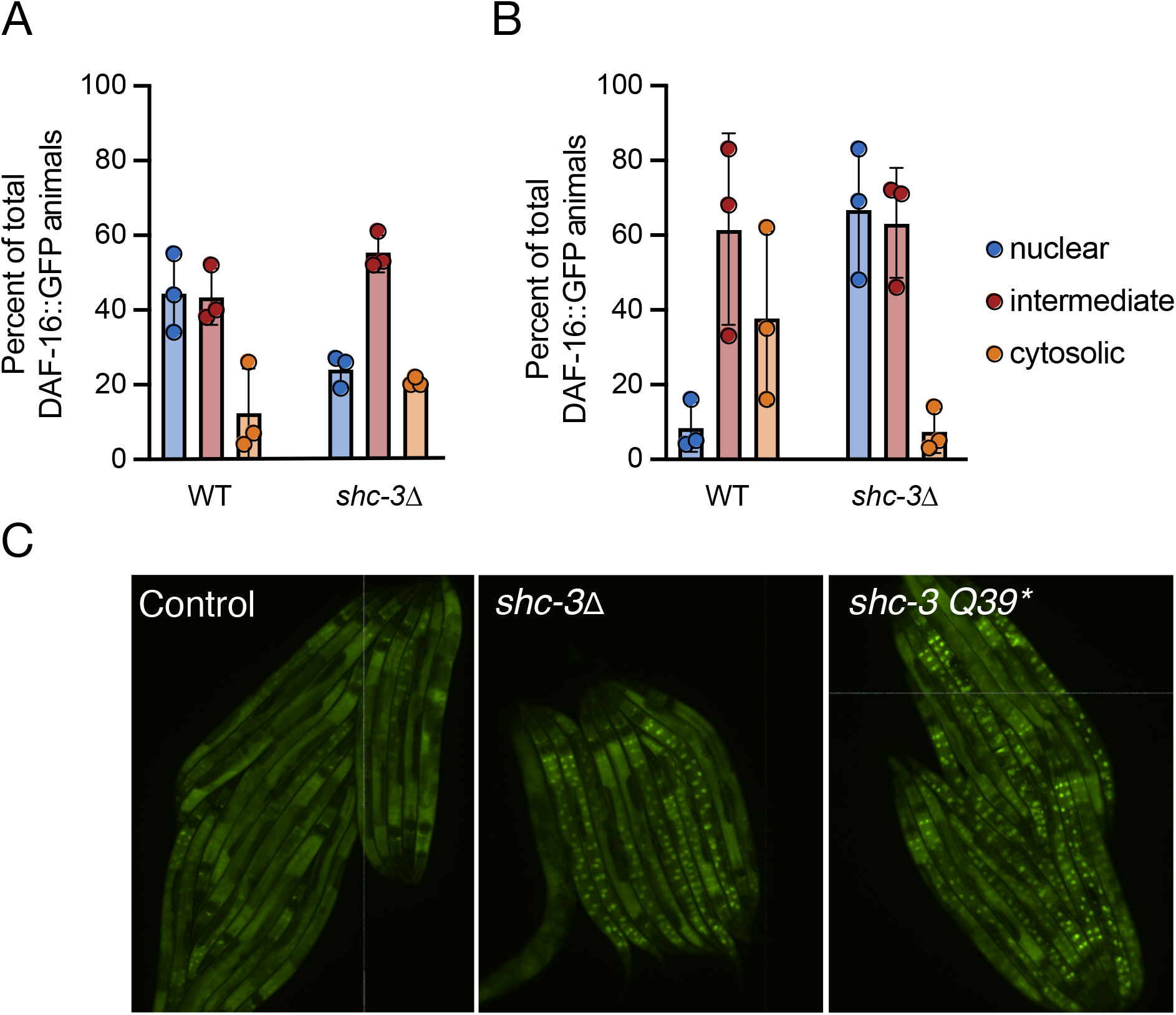
SHC-3 is required for rapid nuclear entry and exit. (A) Animals expressing DAF-16::GFP were categorized as having mostly nuclear, intermediate, or mostly cytoplasmic localization of DAF-16::GFP immediately following a 20 minute heat shock. (B) DAF-16 nuclear exit was measured following a 1hr heat shock and 1.5hr recovery period. Animals were categorized as in A. For A and B, each point represents a population of at least 30 animals. Heat shock was done in triplicate. C. Delayed DAF-16::GFP nuclear exit following recovery from heat shock (as in B).

Based on the requirement of SHC-3 to promote rapid nuclear localization of DAF-16, we examined SHC-3 localization at different temperatures and after heat shock. Intriguingly, at 15°C, SHC-3::GFP was mostly cytoplasmically localized (**Figure 6A**). When plates were shifted from 15°C to 37°C, SHC-3::GFP became membrane-localized, suggesting that SHC-3 functions during heat shock at the cell membrane (**Figure 6B**). Because the PTB domain promotes membrane localization, and DAF-2 contains two NXXY PTB-binding motifs, it is likely that SHC-3 regulates IlS through direct interaction with activated DAF-2. This interaction likely modifies IIS rather than strictly acting as a positive or negative regulator of the canonical signaling pathway. This signaling modification may promote more rapid DAF-16 response to changes in insulin signaling.

**Figure 6.**
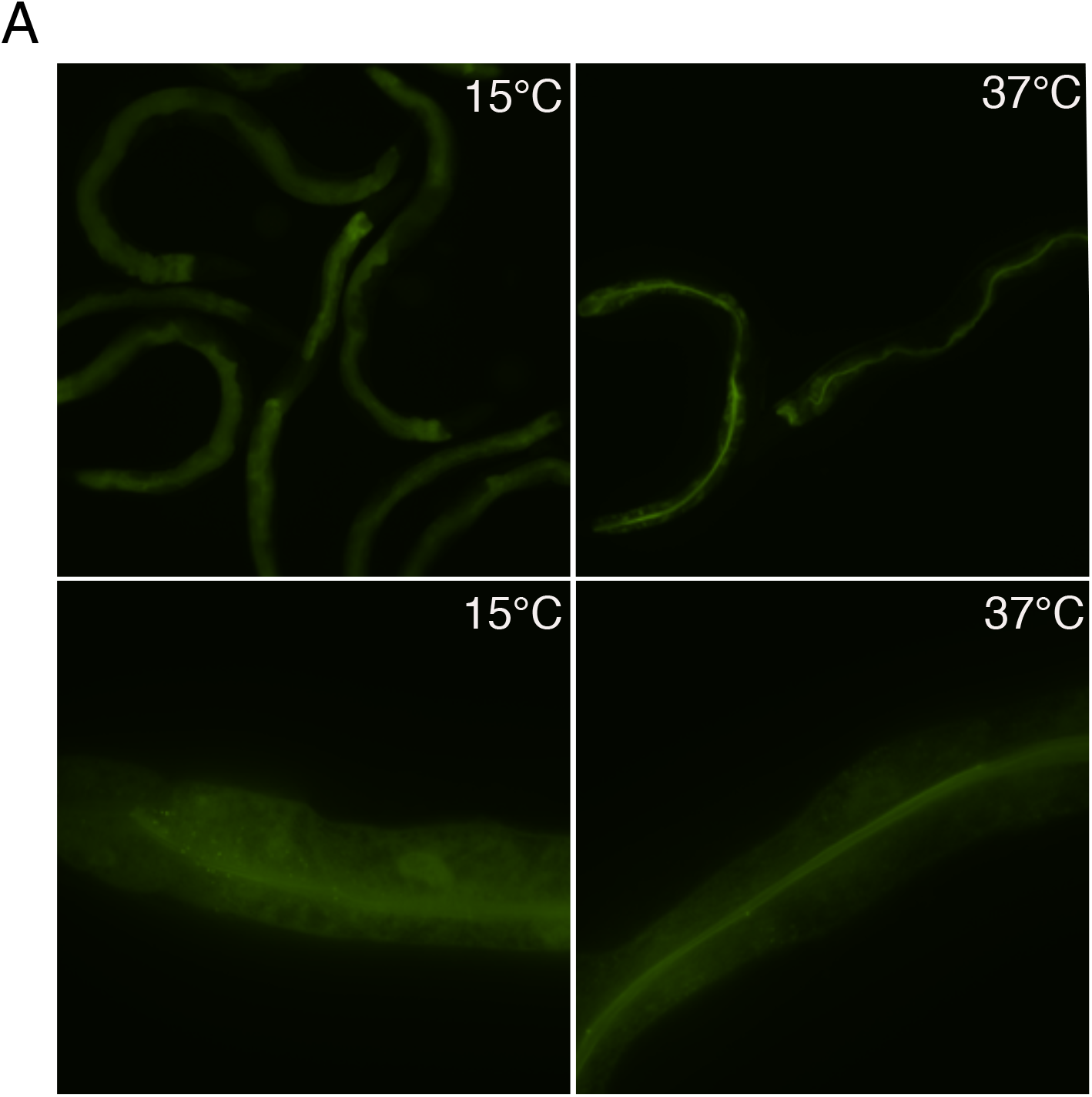
SHC-3 localization is temperature dependent. SHC-3::GFP is found in the cytoplasm when animals are maintained at 15C, but is relocalized to the membrane following heat shock. A. Animals maintained at 15C (left) compared to animals incubated at 37C for 1 hr (right)

## Discussion

Many regulators influence the nuclear localization and activity of DAF-16, together determining the timing and extent of IlS. DAF-16 localization and activity are regulated by post-translational modifications and protein-protein interactions that involve many signaling mediators. We characterized a previously unidentified *C. elegans* Shc gene, *shc-3*, and show that it functions in the IlS pathway to regulate survival in L1 arrest and DAF-16 response to heat stress. The ability of SHC-3 to enhance both nuclear entrance and exit of DAF-16::GFP suggests that the SHC-3 functions to promote rapid changes in signaling.

Unlike other mediators of IlS, SHC-3 is not widely expressed. This, together with the fact that the mutant phenotypes observed are less severe than those of *daf-16* mutants, demonstrates that SHC-3 is not absolutely required for insulin signaling. We observed expression of a SHC-3::GFP reporter only in the intestine. This restricted expression pattern may suggest that insulin signaling in the intestine is regulated or modified by SHC-3 to accommodate gut-specific functions. The ability to promote more rapid changes in DAF-16 activity may be beneficial in the intestine, where exposure to pathogens and sub-optimal foods may first be encountered and require immediate response.

SHC-3 localization during heat stress suggests that SHC-3 functions at the membrane to modify signaling. The cytoplasmic localization observed at 15°C suggests that SHC-3 requires an interaction partner to keep it at the membrane. Mammalian ShcA is recruited to the Insulin-like growth factor receptor through interaction with a conserved phosphotyrosine motif. This motif is conserved in DAF-2 and likely acts to recruit PTB-domain containing proteins, including IRS-1, SHC-1, and SHC-3, upon receptor activation. SHC-1 can interact with DAF-2 but is predominantly found in the nucleus and cytoplasm(Neumann-Haefelin *et al*. 2008), suggesting function at the membrane may be a secondary role. The localization of SHC-3 to the membrane at higher temperatures suggests that it is present at the membrane when DAF-16 is in the nucleus, suggesting SHC-3 negatively regulates the DAF-2-AGE-1-AKT-1 pathway during heat stress. This may occur through competitive binding with other PTB-domain containing proteins that positively affect DAF-2 signaling or by recruiting other signaling interactors that modify DAF-2 signaling.

Although genetic interactions demonstrate that *shc-1* and *shc-3* function in the same pathway, they play different molecular roles. Mutations in either gene reduce survival in L1 arrest without enhancement of mutant phenotypes in double mutants, suggesting that the two proteins cannot substitute for one another. Interpreting the role of SHC-1 in these interactions is challenging due to the two roles it plays upstream of DAF-16. SHC-1 functions as a positive regulator of the MLK-1-MEK-1-KGB-1 pathway. KGB-1 promotes DAF-16 nuclear localization in larvae (Twumasi-Boateng *et al*. 2012). Because *kgb-1* mutants also have decreased survival in L1 arrest, it is likely that SHC-1 function in this pathway is at least partially responsible for the decreased L1 survival observed in *shc-1* mutants. The subcellular localization of SHC-1::GFP, predominantly in the cytoplasm and the nucleus, support the idea that this is the predominant role of SHC-1 in the intestine.

The ability of *shc-3* mutation to suppress oxidative stress sensitivity and shortened lifespan in *shc-1* mutants could result from the two proteins antagonizing one another. This antagonism could be explained if both SHC-1 and SHC-3 bind DAF-2, but subsequently recruit different mediators, a balance of which is required to generate a context appropriate response. This is consistent with the ability of mammalian Shc proteins to bind to the same target proteins with different outcomes (Liu and Meakin 2002; Finetti *et al*. 2009; Patrussi *et al*. 2014). The balance of protein binding in these cases can fine-tune downstream signaling, providing high or low levels of signaling outputs that are context-appropriate. Alternatively, the outcome of *shc-1* and *shc-3* interactions could reflect the balance of positive and negative activities on DAF-16 and may be determined by which function of SHC-1 is involved and in which tissue. KGB-1 regulation of DAF-16 nuclear localization in the intestine can be mediated cell-nonautonomously by expression of KGB-1 in the nervous system (Liu *et al*. 2018). SHC-1 is therefore likely to function in the same capacity. Interpreting interactions between SHC-1 and SHC-3 is further complicated by the ability of KGB-1 to function as a positive regulator of DAF-16 during development but as a negative regulator in adults (Liu *et al*. 2018).

Increased temperature promotes both DAF-16 nuclear import and export (Singh and Aballay 2009). We find that both import and export are slowed in *shc-3* mutants, which may suggest that in the absence of SHC-3, activation of DAF-2 signaling is slowed or less responsive to stimulus. The cumulative effects of SHC-3 loss would therefore depend on the activity of DAF-2 and of other DAF-16 regulators. For example, if we assume that DAF-2 is active during development before reaching the L1 checkpoint, the loss of SHC-3 may reduce survival because DAF-16 entry into the nucleus is delayed upon starvation. Alternatively, if DAF-16 cycles between the nucleus and the cytoplasm, SHC-3 may alter the time it spends in each compartment, which could impact survival.

The ability to regulate both nuclear import and export may function to promote more rapid changes in insulin signaling. There are advantages to tailoring the speed of the DAF-16 response. In some conditions, rapid response may be favorable, for example in response to toxins and other stresses, when immediate mitigation is advantageous. A slow response may be favored in cases where environmental changes fluctuate rapidly. This slowing may act to dampen the response, preventing dramatic changes in signaling until there is a prolonged stimulus.

## Materials and Methods

### *C. elegans* strains

*C. elegans* were propagated by standard methods (Stiernagle 2006). The *shc-3(syb1634)* deletion allele was generated by Suny Biotech by CRISPR/Cas9. This deletion produces a 588 bp deletion that removes exons 1-4. The *shc-3 (gk890887)* mutation was recovered from VC40938 and outcrossed eight times to N2. This mutation produces a stop codon in exon 2 that truncates the protein at amino acid Q39. The *shc-3(gk466377)* allele is a point mutation that generates the R145Q amino acid substitution. Similarly, *shc-3(gk466377)* was recovered from VC40108 and outcrossed eight generations to N2. VC40938, VC40108, *shc-1(ok198), shc-1p::shc-1::GFP* (McKay et al., 2004), CF1038 *daf-16(mu86)*, and TJ356 *daf-16p::daf-16::GFP* (Henderson and Johnson 2001) were obtained from the CGC.

### *shc-3* reporter strain

The translational *shc-3p::shc-3::GFP* fusion was generated by fusing a genomic DNA fragment containing 3.5Kb of sequence 5’ to the *shc-3* start codon, and sequences from exon 1 to the BglII site in exon 4 in-frame with the remaining *shc-3* cDNA. A GA linker (GAGAGAG) was added and both fragments were inserted in frame with GFP into the fire lab vector pPD95_77. This construct was injected into N2 animals at 25 ng/µL and GFP positive animals were selected. Animals were anesthetized with levamisole, mounted on 1% agarose pads and examined on a Nikon Eclipse II microscope. Five independent lines from two separate injections were examined and produced the same expression pattern.

### Pathogen assays

*C. elegans* slow-killing assays were performed as previously described (Tan, Mahajan-Miklos, and Ausubel 1999). Approximately 80-100 early adult animals were transferred to lawns of *P. aeruginosa* and an *E. coli* OP50 control grown on 6 cm Slow killing Media (SKM) plates supplemented with 150 µM FUDR to prevent egg-laying. Plates were incubated at 25°C and scored for live worms daily. Animals unresponsive to touch with a platinum wire were considered dead and removed from the plate. Survival curves were generated using GraphPad Prism Software Version 9.2.0. statistical significance between treatment conditions was determined using the Log-rank (Mantel-Cox) test.

### Survival assays

Eggs were collected by hypochlorite treatment of densely grown, but not starved, populations. Eggs were washed four times in M9 buffer and resuspended at 2 worms/µL in 7 mL M9 buffer at 20°C. Worms were allowed to hatch overnight and hatched L1s were counted the following day. An aliquot was used to count live and dead worms. Each sample was counted daily in triplicate and survival curves were generated in GraphPad Prism. Survival assays were repeated at least three times.

### Stress response

CuSO4 sensitivity was assessed as described (Neumann-Haefelin *et al*. 2008). Briefly, 50-100 eggs were placed on NGM containing 40 µM CuSO4 and incubated at 20°C. The number of adult animals present after four days of growth was determined. Tunicamycin sensitivity was determined in the same way using 1 µg/mL Tunicamycin.

To test oxidative stress sensitivity, animals were allowed to develop until the population was a mix of L4 and young adult animals. Animals were then washed off with M9 and transferred to NGM plates supplemented with 4 mM paraquat and 100 µM FUDR. Animals were gently prodded with a platinum wire, unresponsive worms were scored as dead. Worms that crawled off the plate were censored. Statistical analysis was performed using GraphPad Prism, P-values were determined by using a Mantel-Cox log-rank test. All assays were carried out in triplicate.

To monitor DAF-16::GFP nuclear entry, animals were incubated for 20 minutes at 37C and examined immediately. To examine nuclear export, animals were incubated for 1 hour at 37°C, at which time GFP was nuclear in most, if not all animals, worms were then transferred to 25°C and examined 1.5 hrs after heat shock. Worms were scored as having DAF-16::GFP mostly nuclear, mostly cytoplasmic, or both nuclear and cytoplasmic (Oh *et al*. 2005). All experiments were repeated in at least three independent trials.

## Competing interest statement

The authors declare no competing interests.

## Acknowledgements

Some strains were provided by the CGC, which is funded by NIH Office of Research Infrastructure Programs (P40 OD010440). We gratefully acknowledge the support of the Natural Sciences and Engineering Research Council of Canada (NSERC), [RGPIN-2016-06339].

## Contributions

VLLG, MDB, WRH and LM carried out experiments. All authors read and provided input on the manuscript.

---

**Supplemental Figure S1.**
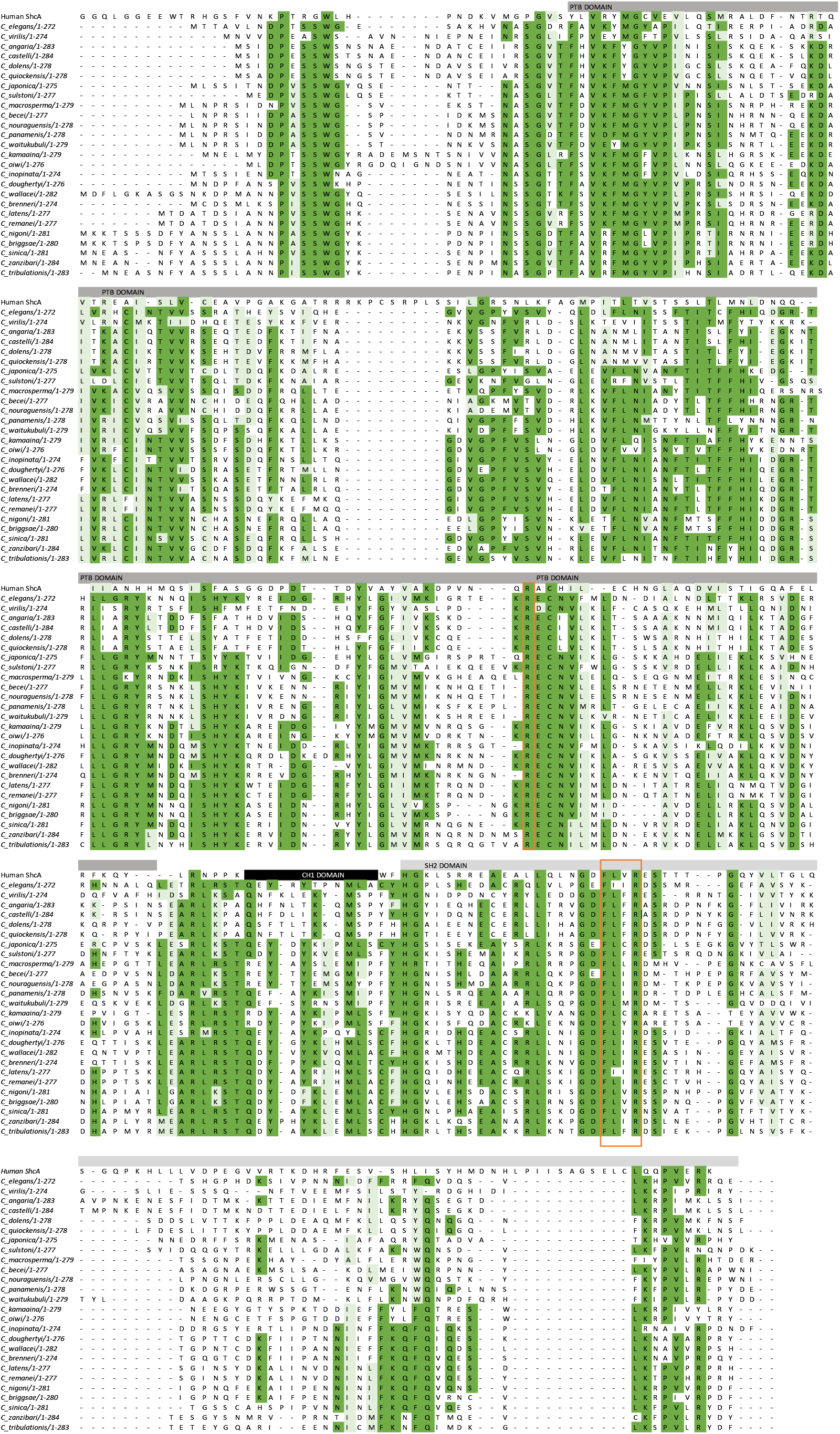
Alignment of Caenorhabitis SHC-3 proteins. Human ShcA is provided for reference (ShcA CH1 domain is not shown). Conserved arginine sequences are boxed in orange.

**Figure S2.**
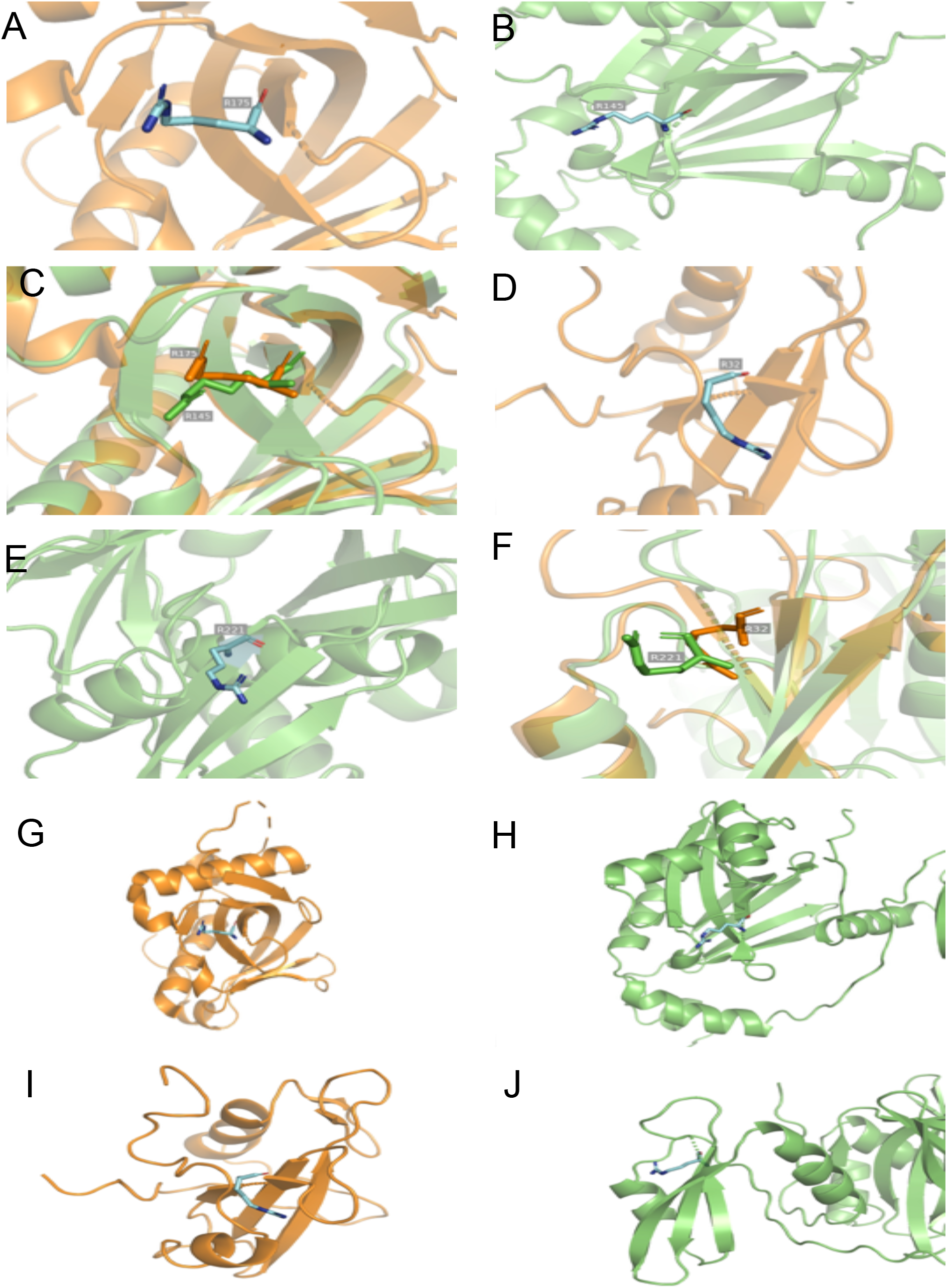
Comparison of mammalian ShcA and C. elegans SHC-3 protein structures. SHC-3 structure from alphafold. a-c PTB domain with conserved Arginine highlighted. d-f SH2 domain with conserved arginine highlighted. Comparison of PTB domains in SHC-3 and mammalian shcA (g-h). Comparison of SH2 domains (i-j)

